# Gain-of-function, focal segmental glomerulosclerosis *Trpc6* mutation minimally affects susceptibility to renal injury in several mouse models

**DOI:** 10.1101/2022.02.11.479954

**Authors:** Brittney J. Brown, Kimber L. Boekell, Brian R. Stotter, Brianna E. Talbot, Johannes S. Schlondorff

**Affiliations:** Division of Nephrology, Beth Israel Deaconess Medical Center, Harvard Medical School, Boston, Massachusetts, USA; Division of Nephrology, Boston Children’s Hospital, Harvard Medical School, Boston, Massachusetts, USA

## Abstract

Mutations in *TRPC6* are a cause of autosomal dominant focal segmental glomerulosclerosis in humans. Many of these mutations are known to have a gain-of-function effect on the non-specific cation channel function of TRPC6. *In vitro* studies have suggested these mutations affect several signaling pathways, but *in vivo* studies have largely compared wild-type and *Trpc6*-deficient rodents. We developed mice carrying a gain-of-function *Trpc6* mutation encoding an E896K amino acid change, corresponding to a known FSGS mutation in *TRPC6*. Homozygous mutant *Trpc6* animals have no appreciable renal pathology, and do not develop albuminuria until very advanced age. The *Trpc6*^*E896K*^ mutation does not impart susceptibility to PAN nephrosis. The animals show a slight delay in recovery from the albumin overload model. In response to chronic angiotensin II infusion, *Trpc6*^*E896K/E896K*^ mice have slightly greater albuminuria initially compared to wild-type animals, an effect that is lost at later time points, and a statistically non-significant trend toward more glomerular injury. This phenotype is nearly opposite to that of *Trpc6*-deficient animals previously described. The *Trpc6* mutation does not appreciably impact renal interstitial fibrosis in response to either angiotensin II infusion, or folate-induced kidney injury. TRPC6 protein and TRPC6-agonist induced calcium influx could not be detected in glomeruli. In sum, these findings suggest that a gain-of-function *Trpc6* mutation confers only a mild susceptibility to glomerular injury in the mouse.

## Introduction

Focal and segmental glomerulosclerosis (FSGS) is a common cause of nephrotic syndrome in adults, frequently progresses to end-stage kidney disease, and has few effective treatment options.[1-3] Studies over the last 25 years have uncovered a substantial genetic component in the pathogenesis of FSGS.[4-9] Several dozen genes are implicated in the development of autosomal recessive and dominant forms of congenital nephrotic syndrome and FSGS.[10] Among these, gain-of-function mutations in *TRPC6* are known to cause autosomal dominant FSGS in humans.[11-17]

Canonical transient receptor potential 6 (TRPC6) is a member of the transient receptor potential (TRP) superfamily of cation channels.[18, 19] TRPC6 is a non-specific cation channel activated downstream of Gα_q_ coupled receptors,[20] including angiotensin II receptor 1,[13] and functions as a receptor-operated calcium effector.[21] Multiple *TRPC6* mutations have been reported in cases of autosomal dominant, predominantly adult onset, FSGS.[12-17, 22] The plurality of these have a gain-of-function (GOF) phenotype, and cluster to the interface of the N-terminal ankyrin repeat domain with the C-terminal rib helix and coiled-coil,[23-25] disrupting an inhibitory calcium-binding site.[26] *In vitro* studies demonstrate that overexpression of TRPC6 GOF mutants activates several signaling pathways,[27, 28] and induces cytotoxicity.[29-31] The relevance of these findings to disease pathogenesis, however, remains uncertain.

An animal model genetically recapitulating TRPC6 gain-of-function disease has not been reported to date. Transgenic overexpression of either wild-type, or gain-of-function, mutant *Trpc6* in podocytes causes only modest albuminuria and mild histological changes in mice, with no clear difference between wild-type and mutant *Trpc6* animals.[32] As *Trpc6* is widely expressed, including in mesangial cells, renal tubular epithelial cells, smooth muscle cells, and fibroblasts,[13, 33-35] the relative importance of TRPC6 activity in podocytes versus other cells types in the pathogenesis of FSGS is unclear. While *TRPC6* has been shown to be upregulated in several acquired proteinuric diseases [36-38], whether increased channel expression mediates similar pathologic effects as mutant TRPC6 is unknown. Studies utilizing *Trpc6* knockout animals have provided conflicting data as to a role for the wild-type channel in renal disease, with some reporting amelioration of disease [29, 35, 37, 39-41], and others increased susceptibility.[42, 43]

In the present study, we characterize the renal consequences of introducing a gain-of-function E896K mutation,[44] corresponding to the human TRPC6 E897K FSGS mutation,[12] into the mouse *Trpc6* gene. Homozygous *Trpc6*^*E896K/E896K*^ mice have no baseline renal pathology or proteinuria until advanced age. They show no susceptibility to PAN nephrosis, only mildly delayed recovery from the albumin overload model, and transiently higher albuminuria early in an angiotensin II infusion model with an associated trend toward more glomerular sclerosis. Furthermore, the *Trpc6* mutation does not influence recovery from, nor residual interstitial fibrosis induced by, folate-induced acute kidney injury. In sum, the results suggest that gain-of-function *Trpc6* mutations induce only mild susceptibility to renal disease in the mouse.

## Material and Methods

### Materials

All chemicals were purchased from Sigma Aldrich unless otherwise specified. GSK1702934A (GSK) was obtained from Focus Biomolecules and dissolved in DMSO. Fura-2 QBT was from Molecular Devices. Puromycin aminonucleoside was from MedChemExpress.

### Mice

All animal procedures were approved by the Beth Israel Deaconess Medical Center (BIDMC) Animal Care and Use Committee, and carried out in accordance with the National Institutes of Health Guide for the Care and Use of Laboratory Animals. *Trpc6*^*E896K/E896K*^ mice were generated at the BIDMC transgenic core via established protocols as described.[44] Animals were backcrossed and maintained on an FVB/NJ (Jackson Laboratory) background. Genotyping of the *Trpc6*^*E896K*^ locus was performed using a custom TaqMan SNP assay. *Trpc6*^*-/-*^ mice [45] were obtained from the Jackson Laboratory. After crossing the mice with C57BL/6J, heterozygous *Trpc6*^*+/-*^ mice were crossed to generate *Trpc6*^*-/-*^ mice and littermate *Trpc6*^*+/+*^ wild-type animals. Animals were maintained in a temperature controlled facility with a 12 hour light, 12 hour dark cycle, and had ad lib access to water and standard chow.

Serum samples were obtained by cheek pouch vein or terminal cardiac puncture, and sent to the UAB O’Brien Center Core C for serum creatinine measurement by isotope dilution LC-MS/MS. Spot urine samples were collected from mice in unlined cages for up to 3 hours. Albuminuria quantification was performed using a mouse albumin ELISA kit (Bethyl Laboratories, Inc. E90-134). Urine creatinine measurement was performed by quantitative colorimetric assay using the QuantiChrom Creatinine Assay Kit (BioAssay Systems DICT-500).

### Albumin overload model

8-12 week old, male wild-type (n=11) and *Trpc6*^*E897K/E897K*^ (n=15) mice were given daily injections of low-endotoxin bovine serum albumin (BSA, A-9430, Sigma Chemical Co, St. Louis, MO) in sterile saline (300mg/ml) by intraperitoneal injection at a dose of 10mg/g body weight on days 1-5 using an established protocol.[46] Urine samples were collected at baseline, day 2 (prior to the second injection), day 6 (24 hours after the last injection), day 9 and day 12. Urine albumin and creatinine measurements were obtained as above.

### Puromycin aminonucleoside nephrosis

We utilized the two dose PAN model as described by Refaeli et al.[47] 8 to 10 week old male mice of various genotypes were utilized: *Trpc6*^*E896K/E896K*^ on an FVB background; and wild type and *Trpc6*^*+/E896K*^ F1 offspring of FVB and 129X1/SvJ (Jackson Laboratory) crossings. Animals were given two doses of puromycin aminonucleoside (450mg/kg; dissolved in normal saline at 18mg/ml) by intraperitoneal injection on days 0 and 7. Urine was collected at baseline and at day 14 and 21; animals were sacrificed and kidneys collected at day 14 or 21.

### Angiotensin II infusion model

Three month old male wild-type and *Trpc6*^*E896K/E896K*^ mice (n=10 each) were implanted with osmotic minipumps (Alzet model 2004; Alza Corp) loaded with angiotensin II diluted in sterile saline to provide a dose of 1 µg/kg/min. The minipumps were implanted in a dorsal subcutaneous location under isofluorane anesthesia under sterile conditions. Urine was collected at baseline, and at weeks 2 and 4 of the infusion. Serum was collected at baseline and at the time of sacrifice. At the end of the 4 week infusion period, animals were sacrificed, and heart and kidneys were harvested and weighed. Tissue samples were processed for histologic analysis. Additional tissue samples were snap frozen in liquid nitrogen for RNA isolation.

### Folate nephropathy model

Three to four month old, male and female, wild-type and *Trpc6*^*E896K/E896K*^ mice (n=13-19 per group) were administered folate (20 mg/ml in 0.3 M sodium bicarbonate solution) at a dose of 250 mg/kg by intraperitoneal injection. Serum samples were collected at baseline (1 week prior to folate administration), 2 days after folate injection, and at the time of sacrifice. 21 days after folate injection, animals were sacrificed under anesthesia. Kidneys were decapsulated and processed for histology and RNA isolation.

### Histology

Tissue was transversely bread loafed and immersion fixed in 4% paraformaldehyde at 4°C for 24 hours. After washing in PBS, samples were further processed for paraffin embedding, sectioning, and staining by the BIDMC Histology Core. Three-micron sections stained with H&E, PAS, and Sirius Red were analyzed using an Olympus BX60 microscope equipped with a digital DP73 camera and cellSens software.

Glomeruli were scored for sclerotic lesions (present or absent) on PAS stained sections by an observer blinded to genotype and treatment. All glomeruli (>100/animal) on a single histologic section containing 2-3 transversely bread loafed portions of a kidney were scored, and the percentage of sclerosed glomeruli calculated.

To calculate podocyte density, kidney sections were stained for immunofluorescence microscopy as described previously,[48] using rabbit monoclonal anti-WT1 (Abcam, ab89901) and appropriate fluorophore-labeled secondary antibody (Jackson ImmunoResearch Laboratories), and counterstained with fluorescently labeled wheat germ agglutinin (Invitrogen, W7024) and Hoechst 33342. Podocyte number and glomerular cross-section were obtained for 20 glomeruli per animal.

Renal fibrosis was quantified by imaging Sirius Red stained sections under polarized light using established methods.[49] Ten non-overlapping 20x fields of cortex were captured per animal, and analyzed using ImageJ. Perivascular areas were excluded from the analysis.

### Platelet and glomerular isolation

Mouse platelets were isolated using modified standard procedures,[50] as previously outlined.[44] Washed platelets were resuspended directly in NP-40 lysis buffer containing Complete protease inhibitors (Roche). Glomeruli were isolated using magnetic bead perfusion as previously described,[48, 51] utilizing high iron content magnetic particles (AMS-40-10H, Spherotech). For western blot, glomeruli were resuspended in NP-40 lysis buffer with protease inhibitors. For podocyte outgrowth, glomeruli were cultured in 6 well tissue culture plates with RPMI-1640 supplemented with 10% fetal bovine serum, and 1% ampicillin, penicillin, and streptomycin.[52] After 10 days of culture, outgrown podocytes were passaged and plated onto clear bottom 96 well plates for calcium imaging experiments.

### Fura-2 calcium imaging

Primary podocyte cultures were subject to Fura-2 fluorescence ratio measurement (Fura-2 QBT kit R8197, Molecular Devices) using a FlexStation III reader with automated pipetting at the Harvard ICCB-Longwood screening facility in a 96-well format essentially as previously described [30, 44]. Final concentrations of agonists were as follows: 100 µM ADP, 100 µM ATP, 50 µM GSK1702934A, 0.5 u/ml thrombin, and 1 µM thapsigargin.

### RNA isolation and gene expression

RNA was isolated from snap frozen tissue using RNeasy universal kits, and reverse transcribed into cDNA with the QuantiTect Reverse transcription kit (both Qiagen). Real-time PCR reactions were run on a QuantStudio 6 Flex machine using PowerUp SYBR Green master mix (Applied Biosystems) using gene specific primer sets (Table 1). Expression levels were normalized to 18S rRNA and *Gapdh*.

**Table 1.**
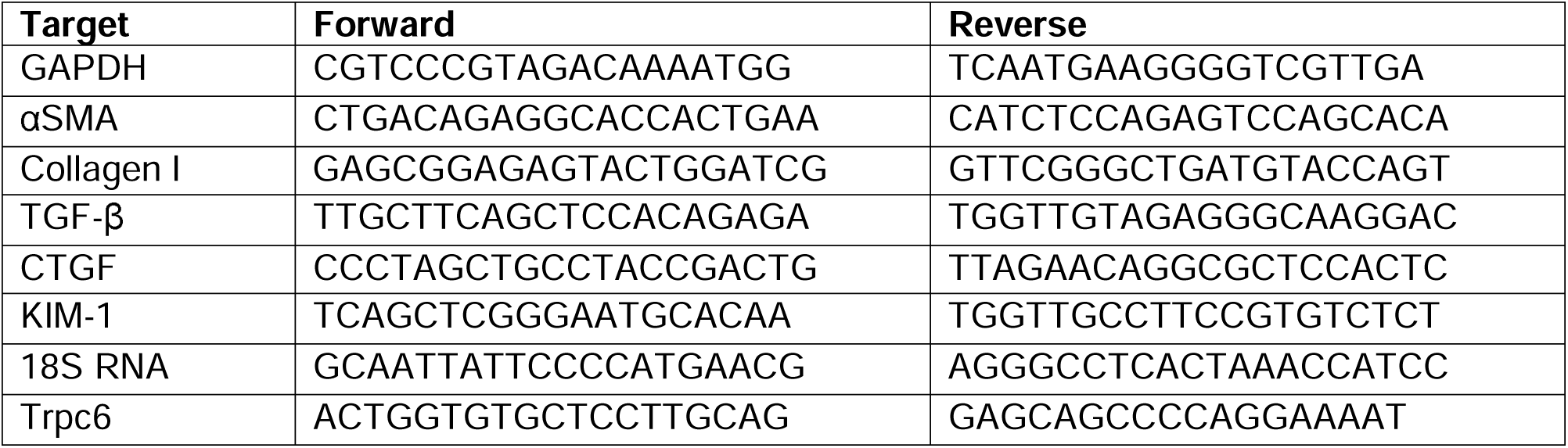
Primer sequences used for qPCR gene expression analysis.

### Western blotting

Lysates were mixed with 4x sample loading buffer containing β-mercaptoethanol and immediately incubated at 95°C for 5 minutes. SDS-PAGE was performed as previously described.[28] Western blotting using fluorescence detection was performed using Immobilon-FL PVDF membrane (Millipore), Chameleon Duo pre-stained protein ladder, Intercept blocking buffer, and an Odyssey CLx imaging system (all LI-COR). Primary antibodies against the following antigens were utilized: Erk1/2 (CST #9107, 1:1000), Podocalyxin (MAB1556, R&D Systems, 1:500), TRPC6 (Alomone ACC-017, 1:500). Fluorescent secondary antibodies (IRDye 690RD anti-mouse and anti-rat; IRDye 800CW anti-rabbit; all LI-COR) were used at 1:20,000 dilution.

### Statistical analysis

All statistical analyses were performed using GraphPad Prism version 9. The specific statistical tests utilized for each experiment are specified within the corresponding figure legends. Symbols used for pair-wise comparison adjusted p-values are: ns, p>0.05; *, p<0.05; **, p<0.01; ***, p<0.001; ****, p<0.0001.

## Results

*Trpc6*^*E896K/E896K*^ mice were viable and fertile, and born at the expected Mendelian ratio when generated by mating heterozygous animals. *Trpc6*^*+/E896K*^ and *Trpc6*^*E896K/E896K*^ mice showed no evidence of developing albuminuria compared to their wild-type counterparts at 6 months of age (Fig. 1A). Even in female animals aged 20 to 23 months, albuminuria did not differ significantly between wild-type and knock-in animals (Fig. 1B). The fraction of *Trpc6*^*E896K/E896K*^ mice that did develop albuminuria demonstrated mesangial expansion and mesangial hypercellularity, but no appreciable glomerular sclerosis (Fig. 1C), similar to age-associated glomerular changes reported to occur in wild-type female mice in this age range.[53, 54]

**Fig 1.**
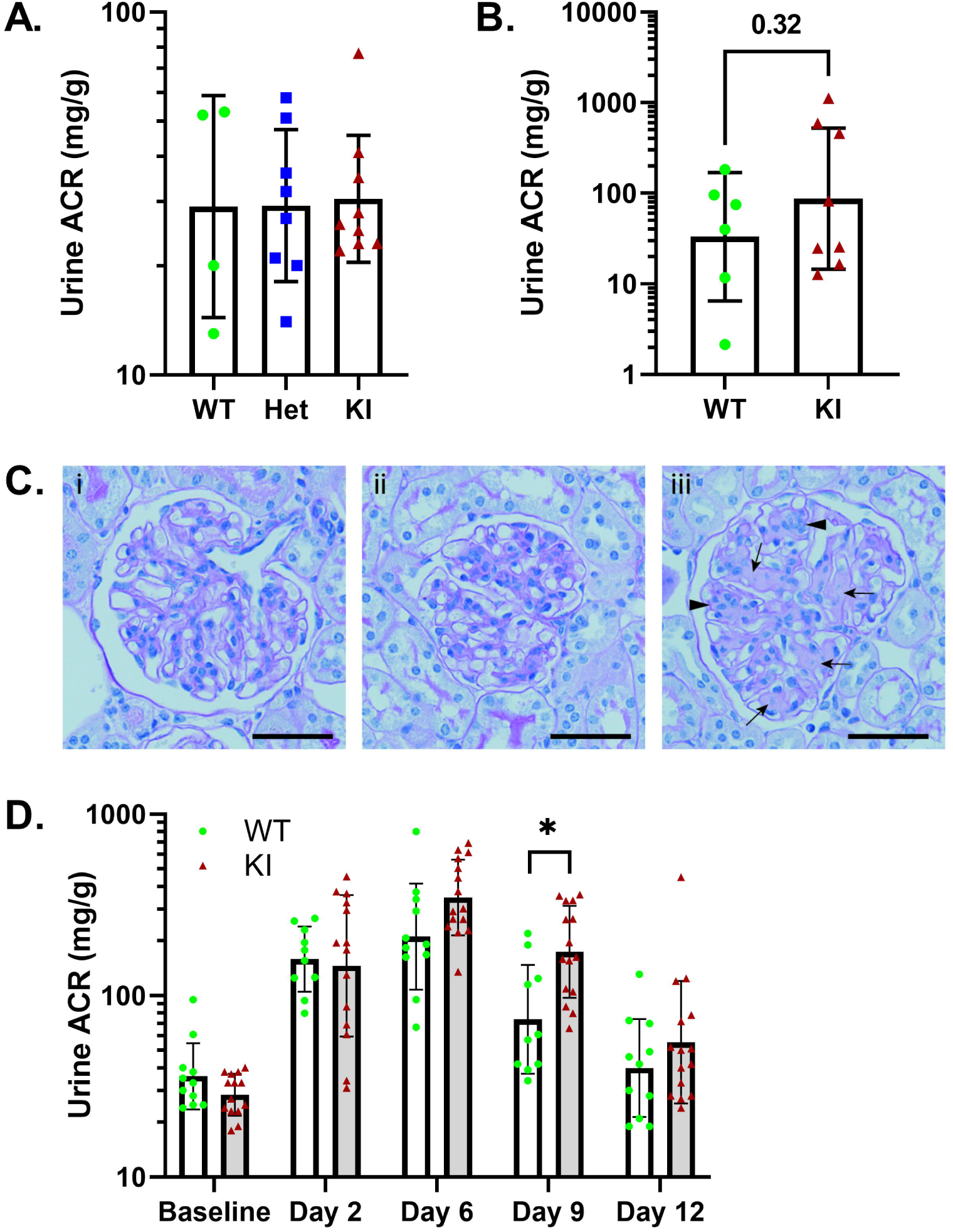
Baseline and albumin overload induced albuminuria in wild-type and *Trpc6*^*E896K/E896K*^ mice. A, urine albumin-to-creatinine ratio (ACR) measurements from six-month old, wild-type (WT), *Trpc6*^*+/E896K*^ (Het), and *Trpc6*^*E896K/E896K*^ (KI) males. Shown are individual values, geometric mean and SD; n=4-9/group. Groups were compared by one-way ANOVA with Tukey’s multiple comparisons test; all comparisons without statistical significant differences. B, urine ACR from 20-23 month old, female WT (n=6) and KI (n=8) animals. Shown are geometric mean and individual values; log-transformed ACRs were compared by unpaired t-test. C, glomerular histology of 23 month old, female (i) wild-type, (ii) non-proteinuric KI, and (iii) albuminuric KI mice. Mesangial expansion (arrows) and mesangial hypercellularity (arrowheads) are apparent in the albuminuric animal. PAS stained sections; scale bar represents 20 µm. D, albuminuria in WT and KI male mice subject to albumin overload from day 1-5. Shown are geometric mean, SD and individual values; n=11 (WT) and 15 (KI). Log-transformed ACRs were compared between genotypes by mixed-effects analysis with Sidak’s multiple comparisons test. *Trpc6*^*E896K/E896K*^ had statistically significantly more albuminuria compared to wild-type only on day 9.

### Albumin overload model

Wild-type and *Trpc6*^*E896K/E896K*^ male mice were compared in their response to the transient albumin overload model (Fig. 1D). A mixed-effects model suggests a statistically significant effect of genotype on albuminuria across the combined time-points (p=0.0153). However, by multiple comparisons, the albuminuria observed in *Trpc6*^*E896K/E896K*^ male mice was only statistically significantly higher than controls at day 9, 4 days after the last albumin administration. Both groups showed a similar degree of albuminuria a week after injections were discontinued, when albuminuria had returned back to near baseline. As has been reported by others,[55] no histological sequelae were apparent by light microscopy after the recovery period.

### Puromycin aminonucleoside nephrosis

Mice heterozygous for the *Inf2*^*R218Q*^ mutation,[56] or a *Podxl* null mutation,[47] show dramatic sensitivity to glomerular injury in the PAN model. We therefore examined whether the *Trpc6*^*E896K*^ allele might confer a similar susceptibility. *Trpc6*^*E896K/E896K*^ mice exposed to the two dose PAN regimen[47] had only a small increase in proteinuria (Fig. 2A) without the development of glomerular sclerosis (Fig. 2B). We also compared *Trpc6*^*+/+*^ and *Trpc6*^*+/E896K*^ animals generated through an F1 cross with Sv/129 animals, as Sv/129 animals show some susceptibility to PAN (data not shown). These animals all developed low grade proteinuria (Fig. 2C), but histological analysis revealed no glomerular sclerosis, and only very rare evidence of tubular protein reabsorption droplets and proteinaceous casts (Fig. 2D). Furthermore, albuminuria did not differ based on *Trpc6* genotype. These results suggest that the *Trpc6*^*E896K*^ allele does not affect susceptibility to PAN.

**Fig 2.**
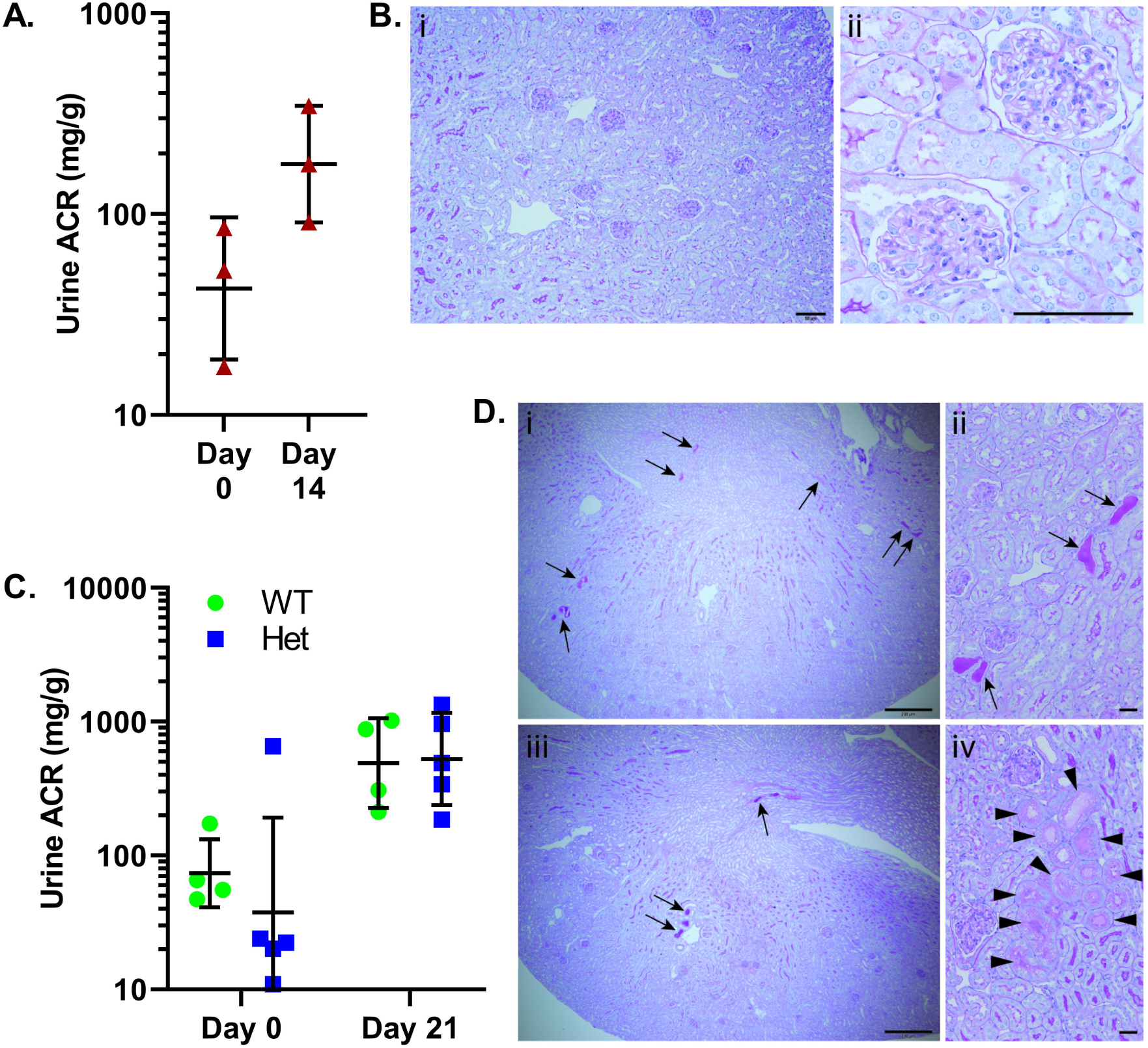
Puromycin aminonucleoside nephrosis in *Trpc6*^*E896K*^ mutant mice. Various *Trpc6* genotype male mice were subject to two intraperitoneal injections of puromycin aminonucleoside (PA; 450mg/kg) on days 0 and 7. *Trpc6*^*E896K/E896K*^ mice on the FVB background developed a small (<5 fold) increase in albuminuria at day 14 compared to baseline (A). B, PAS stained histology sections of *Trpc6*^*E896K/E896K*^ kidney 14 days after PAN revealed no appreciable (i) cortical, or (ii) glomerular, pathology. Scale bar represents 50 µm. C, urinary albumin-to-creatinine ratios (ACR) of *Trpc6*^*+/+*^ (WT) and *Trpc6*^*+/E896K*^ (Het) animals on an F1 FVB/NJ x Sv/129 background before and 21 days after PAN induction. D, PAS stained WT (i, ii), and Het (iii, iv) kidney sections 21 days after initial PA administration demonstrate largely preserved cortical, and glomerular architecture, with very rare (estimated <1% of renal cortex cross sectional area) proteinaceous casts (arrows), and tubules with protein reabsorption droplets (arrowheads). Scale bars represent 200 µm (i, iii), and 20 µm (ii, iv). ACRs (A, C) are shown as geometric mean, SD and individual values. Log-transformed ACRs were compared for statistical analyses by paired t-test or two-way ANOVA.

### Angiotensin II infusion

Multiple studies have reported a role for TRPC6 channel activation downstream of angiotensin II signaling.[13, 57-61] *Trpc6* knockout mice develop less albuminuria initially, and trend toward less renal pathology, compared to wild-type animals, in response to chronic ATII infusion despite a similar response in blood pressure.[39] We therefore exposed *Trpc6*^*E896K/E896K*^ and wild-type male mice to 4 weeks of ATII infusion, and compared their response. *Trpc6*^*E896K/E896K*^ animals developed greater albuminuria after 2 weeks compared to wild-type animals, but this difference did not persist at the end of the infusion period (Fig. 3A). Serum creatinine did not differ between genotypes either at baseline, or at the end of the experiment (Fig. 3B).

**Fig 3.**
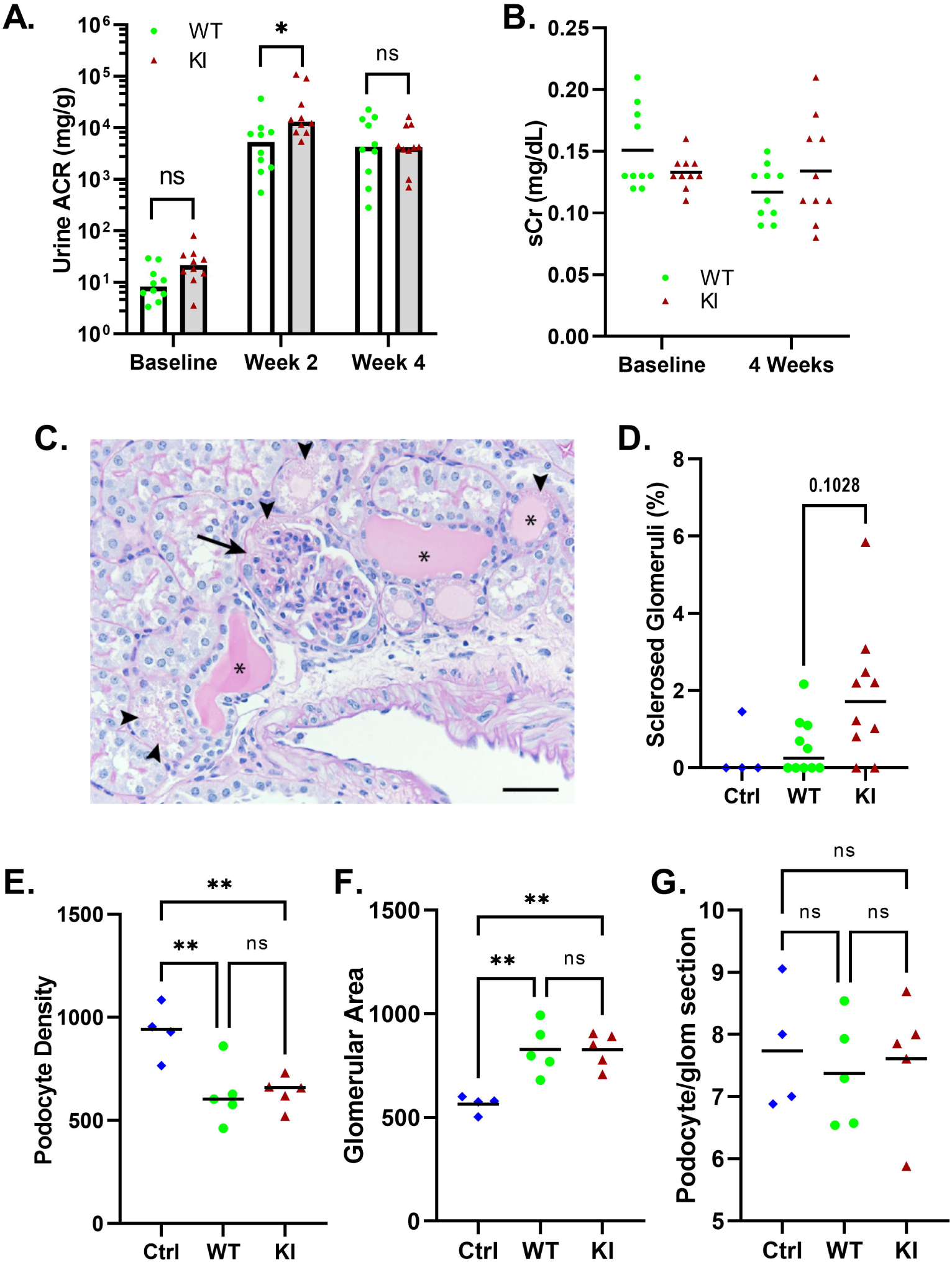
Effect of *Trpc6* genotype on glomerular response to angiotensin II infusion. Wild-type and *Trpc6*^*E896K/E896K*^ (KI) male mice were subject to ATII infusion for 4 weeks. A, urine albumin-to-creatinine ratio (ACR) measurements demonstrate development of robust albuminuria in both groups. KI mice developed slightly greater albuminuria at 2 weeks compared to wild-type, an effect that did not persist at 4 weeks. Shown are median and individual values; n=10/group. Log-transformed ACRs were compared between genotypes by two-way ANOVA with Sidak’s multiple comparisons test. B, serum creatinine measurements at baseline and after 4 weeks of ATII infusion. Shown are mean and individual values; n=10/group; no statistically significant differences between genotypes. C, example of renal pathology in a KI mouse after ATII infusion. Shown is a PAS stained section demonstrating a segmental glomerular lesion (arrow), tubular atrophy with cystic changes and proteinaceous casts (asterisks), and protein reabsorption droplets (arrowheads). Scale bar represents 20 µm. D, percentage of glomeruli showing evidence of segmental lesions in control wild-type males (Ctrl; n=4), and wild-type (WT) and *Trpc6*^*E896K/E896K*^ (KI) males subjected to ATII infusion (n=10 each). Shown are median and individual values; differences between groups did not reach statistical significance by Kruskal-Wallis test with Dunn’s multiple comparisons. Glomerular podocyte density (E), glomerular area (F), and podocyte number per glomerular cross-section (G) were measured. Shown are mean and individual averages per animal in untreated control animals (Ctrl; n=4) and ATII treated wild-type and knock-in animals (n=5 each). 20 glomeruli were measured per animal. One-way ANOVA analysis with Tukey’s multiple comparisons test.

Histologic examination of kidney sections revealed rare glomerular lesions and isolated tubular dilations with proteinacious casts (Fig. 3C). Although there was a trend toward more glomerular lesions in *Trpc6*^*E896K/E896K*^ mice, this did not reach statistical significance (Fig. 3D). Podocyte density was lower in angiotensin II treated animals compared to controls (Fig. 3E), but *Trpc6* genotype did not affect this parameter. The effect was driven by an increase in average glomerular cross-sectional area (Fig. 3F), with no evidence of significant podocyte loss (Fig. 3G).

Angiotensin II treated animals developed significant perivascular fibrosis, especially in the heart (Fig. 4A). However, interstitial fibrosis of the renal cortex, quantified by birefringence of Sirius red stained sections imaged under polarized light, showed no difference between control and ATII treated animals (Fig. 4B). Kidney (Fig. 4C) and heart (Fig. 4D) weight, normalized to body weight, similarly were not different between groups. *Trpc6* and collagen I gene expression were ascertained by RT-PCR of whole kidney RNA. While expression of both genes was higher in angiotensin II exposed kidneys compared to wild-type control samples, *Trpc6* genotype did not influence the expression levels of the mRNAs after angiotensin treatment (Fig. 4E).

**Fig 4.**
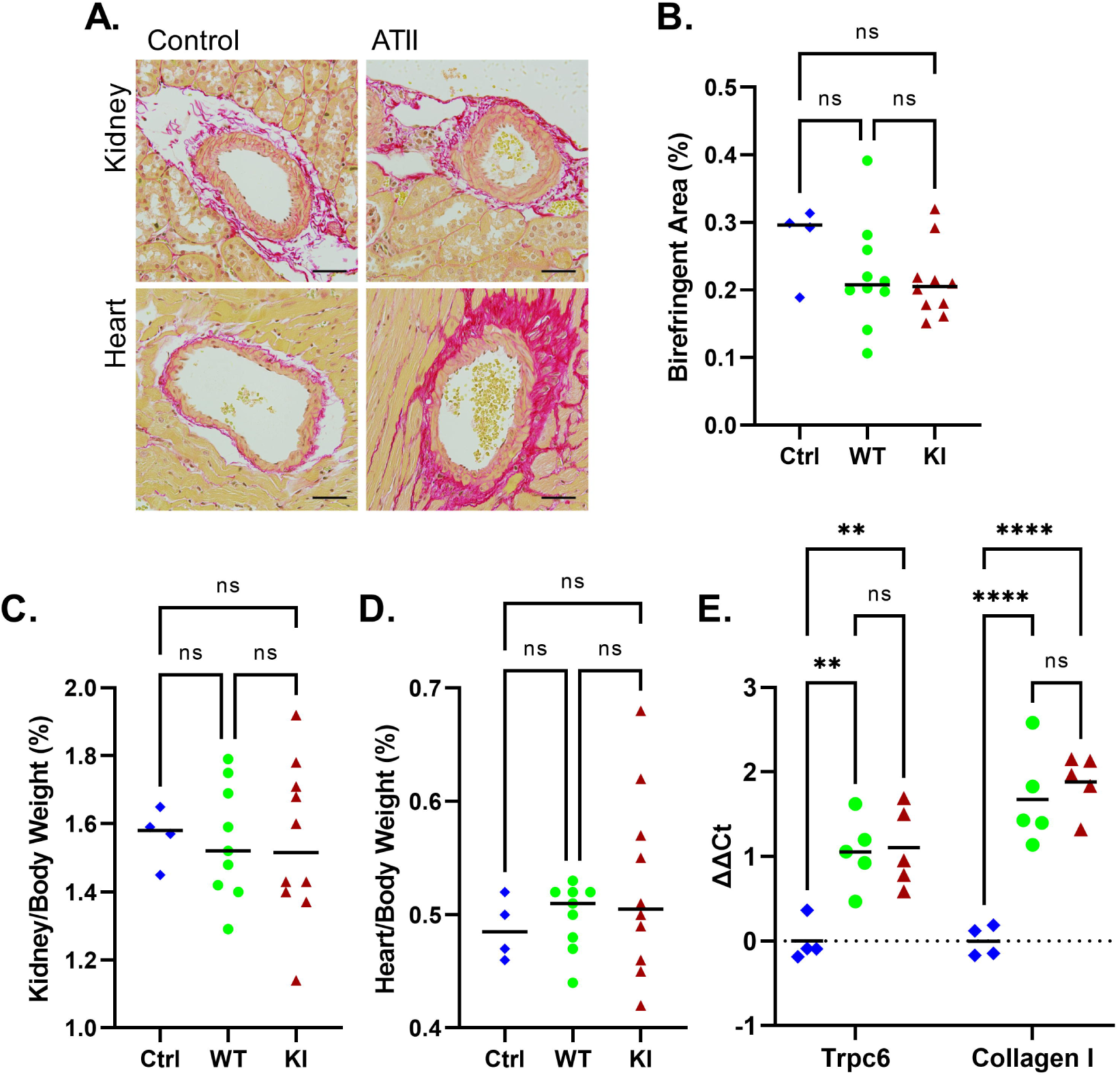
Fibrotic response to angiotensin II infusion. A, histology of control and ATII treated kidneys and heart demonstrating the development of perivascular fibrosis. Sections stained with Sirius Red; scale bar equals 20 µm. B, percentage of Sirius Red stained renal cortex demonstrating birefringence in control wild-type male animals (Ctrl), and wild-type and *Trpc6*^*E896K/E896K*^ (KI) males subjected to ATII infusion. Shown are median and individual values; one-way ANOVA with Tukey’s multiple comparisons. Kidney (C) and heart (D) weights, normalized to total body weight, did not differ between groups. Shown are median and individual values (n=4-10/group); one-way ANOVA with Tukey’s multiple comparisons test. E, relative gene expression analysis demonstrated upregulation of both *Trpc6* and Collagen I mRNA in kidneys subjected to ATII infusion. Shown are mean and individual values; two-way ANOVA with Tukey’s multiple comparisons.

### Folate nephropathy

The administration of a high dose of folic acid leads to tubular injury and AKI in mice.[62-64] Although renal excretory function largely recovers, residual interstitial fibrosis and other hallmarks of chronic injury remain. As TRPC6 has been implicated in modulating fibrosis,[35, 65-69] we compared *Trpc6*^*E896K/E896K*^ and wild-type mice in the folate nephropathy model (Fig. 5). Serum creatinine in both male (Fig. 5A) and female (Fig. 5B) mice demonstrated a robust rise two days after folate administration, returning to baseline after 3 weeks. Serum Cr did not differ significantly between *Trpc6* genotypes at any of the time points. Histologic analysis revealed foci of interstitial fibrosis and tubular atrophy (Fig. 5C). The percentage of cortex demonstrating fibrosis showed significant variability within each group, and no significant difference between *Trpc6* genotypes (Fig. 5D). Similarly, gene expression analysis of several fibrosis and kidney injury marker genes did not reveal any significant differences between male wild-type and *Trpc6*^*E897K/E897K*^ kidneys (Fig. 5E). The results do suggest significant inter-individual variability in the degree of renal scarring in this model.

**Fig 5.**
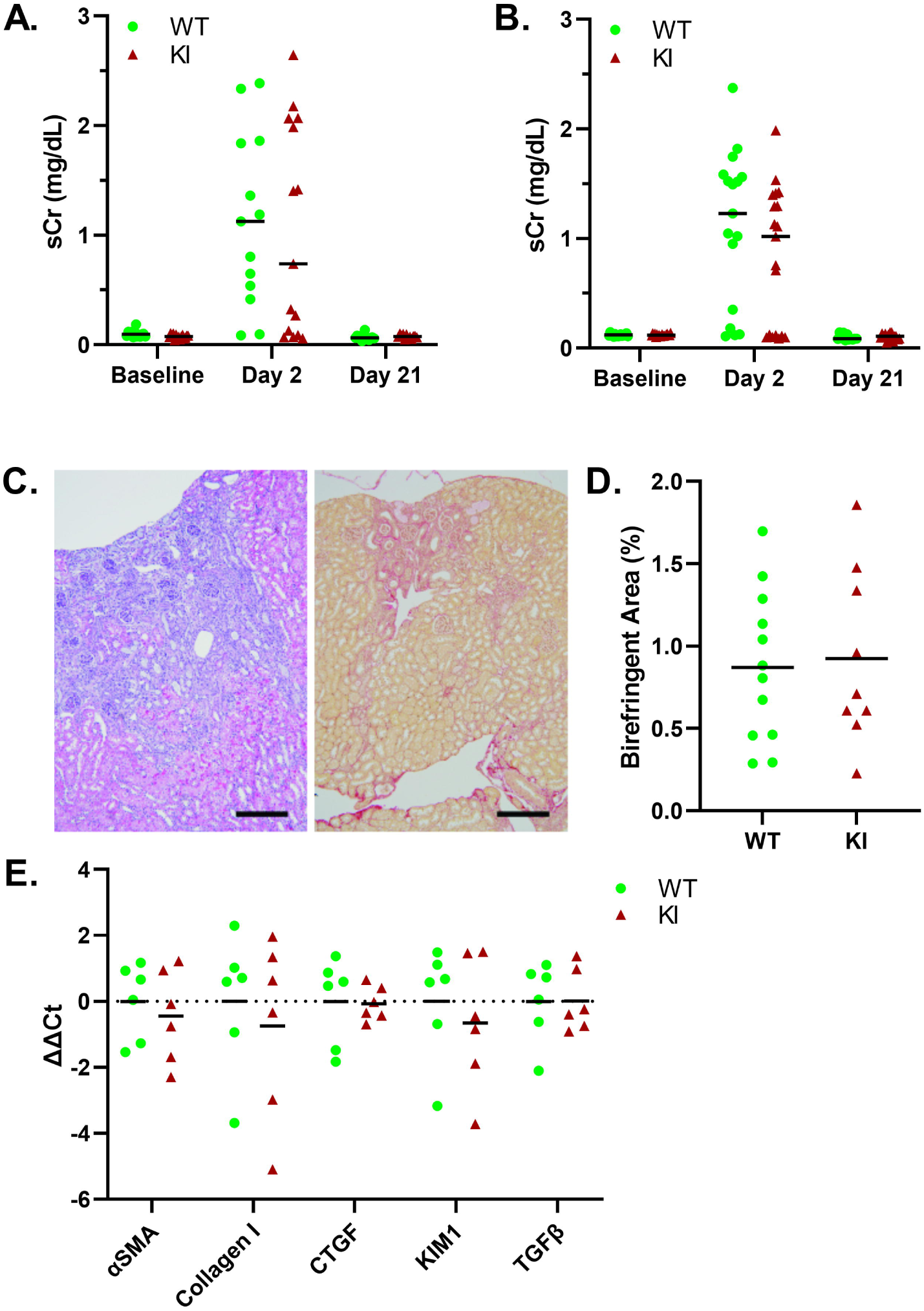
*Trpc6* genotype does not influence response to folate nephropathy. Serum creatinine in WT and KI, male (A) and female (B), mice was elevated 2 days after folate administration, and recovered to baseline levels after 3 weeks. Shown are median and individual values; n=13-19/group. There were no statistically significant differences between genotypes at any of the time-points; two-way ANOVA with Sidak’s multiple comparisons test. C, renal histology at day 21, demonstrating areas of tubular atrophy and interstitial fibrosis surrounded by relatively preserved cortical architecture. Shown are H&E stain of a male wild-type kidney section (left), and Sirius Red stain of a female KI kidney section (right); scale bar equals 100 µm. D, the percentage of Sirius Red stained renal cortex area demonstrating birefringence 21 days after folate-induced AKI was compared between wild-type (n=12) and KI (n=9) male mice. Shown are mean and individual values; no differences between groups by unpaired t-test. E, relative mRNA expression levels of fibrosis and renal injury related genes was compared in WT and KI male folate-nephropathy kidney samples (n=6/group). There were no statistically significant differences between genotypes for any of the genes; multiple unpaired t-tests.

### Glomeruluar TRPC6 expression and function

Although early reports localized human TRPC6 to the glomerular podocyte,[12, 13] *in situ* hybridization,[33, 35] and single cell RNAseq,[70] have not detected substantial *Trpc6* mRNA in murine podocytes. We probed murine glomerular lysates for TRPC6 protein by western blot (Fig. 6A), utilizing *Trpc6*^*-/-*^ mouse samples as negative controls. Although TRPC6 is readily detectable in murine platelets, glomerular extracts show no specific signal. To ascertain if TRPC6-dependent calcium influxes might be detectable despite the lack of TRPC6 signal by western blot, we performed Fura-2 fluorimetry on primary podocytes isolated from *Trpc6*^*E896K/E896K*^ glomeruli (Fig. 6B-D). We utilized GSK1702934A (GSK), a TRPC3/6 agonist [71, 72], as it induces calcium influx in mouse platelets in a TRPC6-dependent manner, with *Trpc6*^*E896K/E896K*^ platelets demonstrating a significantly larger response compared to wild-type platelets [44]. GSK failed to induce any significant change in Fura-2 fluorescence compared to vehicle control. ADP, ATP, and thrombin, utilized as positive controls, all induced Fura-2 responses in these cells, consistent with prior reports [73-77]. In sum, we have been unable to demonstrate the presence of TRPC6 protein, by western blot, or GSK-induced calcium influx in murine glomeruli. We cannot exclude the possibility of low levels of TRPC6 channel, which are not responsive to GSK, being present.

**Fig 6.**
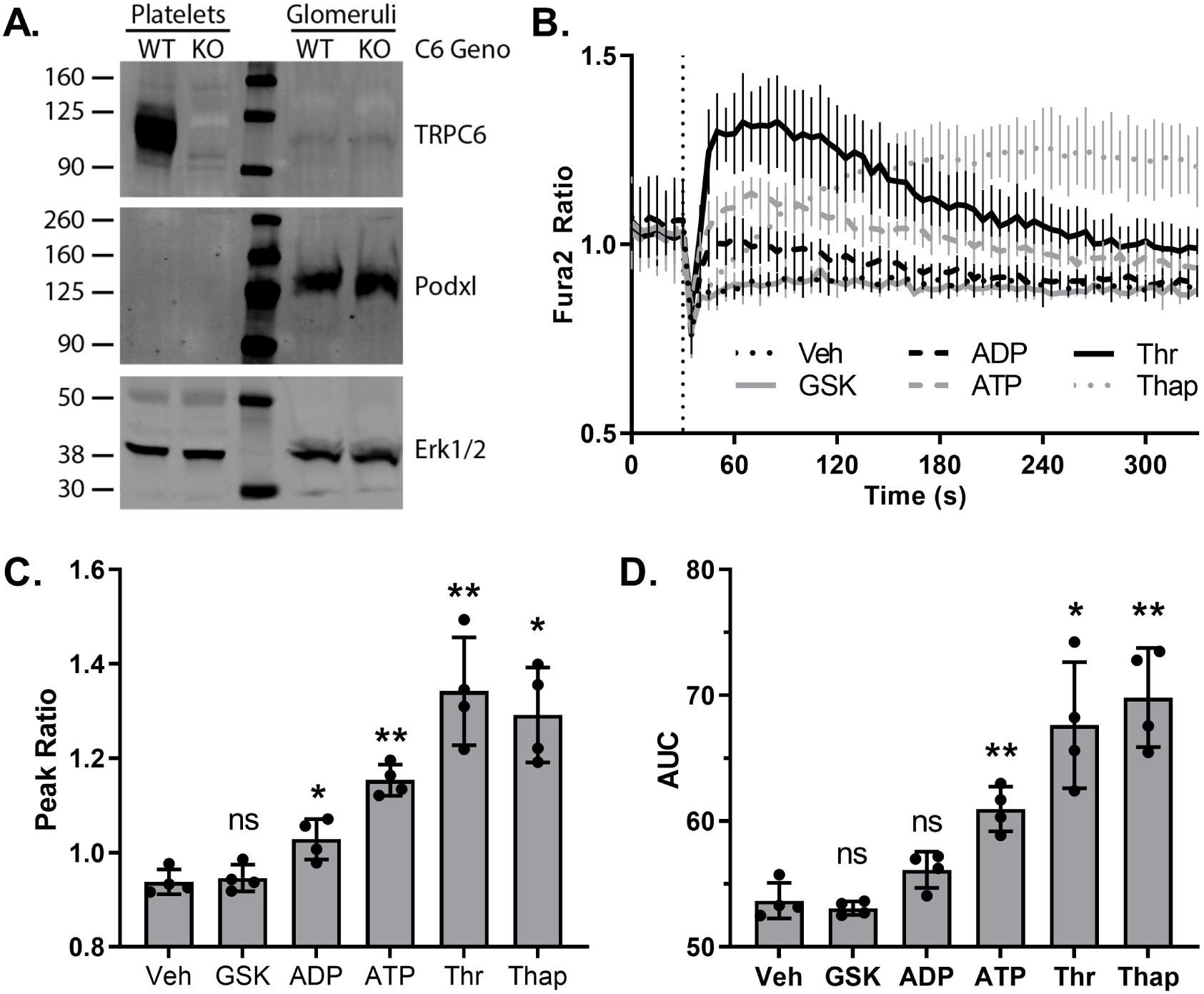
Characterization of TRPC6 expression and GSK1702934A response in murine podocytes. A, platelet and glomerular lysates from wild-type (WT) and *Trpc6*^*-/-*^ (KO) female animals were analyzed by SDS-PAGE and Western blot. Anti-TRPC6 antibodies (top) detected a major band around 105 kD in WT, but not KO, platelets. No corresponding specific band is seen in glomerular lysates. A faint band of a size similar to TRPC6 is seen in both WT and KO glomerular samples, suggesting a non-specific signal. Podocalyxin (Podxl, middle) and Erk1/2 (bottom) blots are shown as loading controls. B, time-course of Fura-2 fluorescence ratio (340/380) in primary podocytes cultured from *Trpc6*^*E896K/E896K*^ (KI) glomeruli. After 30 seconds (dashed vertical line), cells were stimulated with vehicle (Veh), GSK1702934A (GSK; 50 µM), ADP (100 µM), ATP (100 µM), thrombin (Thr; 0.5 u/ml), or thapsigargin (Thap; 1 µM). Shown are mean ± SD; n=4 using podocytes from 2 different animals. Post-stimulation (C) peak Fura-2 fluorescence ratio, and (D) area under the curve (AUC, arbitrary units) were calculated. Shown are the mean and values from individual experiments. RM one-way ANOVA with Dunnett’s multiple comparisons test to vehicle control.

## Discussion

*TRPC6* mutations are a cause of autosomal dominant FSGS in humans.[12, 13] The plurality of these mutations lead to a gain-of-function phenotype at the level of the channel.[12-17, 22] In the current study, we report on the renal phenotype of mice carrying a mutation encoding for a gain-of-function alteration in the TRPC6 protein. Even in the homozygous state and at advanced age, these animals show minimal renal pathology relative to their wild-type counterparts, and no segmental glomerular sclerosis. Furthermore, compared to wild-type animals, they display only mild or no significant differences in their response to several renal stress models. Specifically, they demonstrate slightly delayed resolution of albuminuria in the albumin overload model, only transiently higher albuminuria and a trend toward more glomerular lesions in response to angiotensin II infusion, no enhanced susceptibility to PAN, and no significantly different response to folate-induced AKI and subsequent fibrosis. In sum, these findings suggest that the *Trpc6*^*E896K/E896K*^ mutant mouse does not readily phenocopy the renal pathology associated with gain-of-function *TRPC6* mutations in humans.

Angiotensin II signaling through AT1 receptor is thought to be a central driver in the development of proteinuria and renal injury.[78] *In vitro*, TRPC6 has been shown to be activated downstream of Gα_q_-coupled receptors,[20] and AT1 receptor in particular.[13, 57-61] In a prior study, *Trpc6*^*-/-*^ mice demonstrate slightly lower albuminuria compared to wild-type animals after 2 weeks of ATII infusion, a difference that is no longer significant after 4 weeks; and a non-statistically significant trend toward less glomerular injury.[39] The results presented here, comparing wild-type and *Trpc6*^*E896K/E896K*^ murine responses to ATII infusion, neatly complement these previous results in *Trpc6* knockout animals. *Trpc6*^*E896K/E896K*^ mice show an increase in proteinuria compared to wild-type animals 2 weeks, but not 4 weeks, after the initiation of ATII treatment. Furthermore, there was a trend toward more glomerular lesions in the mutant mice, though the difference did not meet statistical significance. We did not perform blood pressure studies, and so are unable to address whether, as the earlier study[39] did, there was no effect of *Trpc6* genotype on the development of hypertension. Taken together, these results suggest that TRPC6 channel activity contributes, at least transiently, to the development of proteinuria and glomerular lesions in response to high levels of angiotensin II.

TRPC6 has been implicated in the development of fibrosis in several organs.[35, 65-69, 79] In the kidney, both genetic,[35, 67] and pharmacologic,[66] inhibition of TRPC6 is reported to dampen the fibrotic response in the unilateral ureteral obstruction model. We were therefore somewhat surprised that the *Trpc6*^*E896K/E896K*^ mice do not show an increased fibrotic reaction after folate-induced AKI. It is possible that mechanistically, the effects of the gain-of-function *Trpc6* mutation on kidney function are not simply the opposite of deleting *Trpc6*. This has been our experience in platelets.[44] Alternatively, TRPC6-dependent fibrosis pathways may be activated specifically downstream of ureteral obstruction, and not involved in the models tested here. Consistent with this possibility, *Trpc6* knockout in rats does not alter the age-related development of tubulointerstitial fibrosis.[80] Future studies in other fibrosis models will be needed to address these questions.

Genetically recapitulating autosomal dominant forms of FSGS in mice has frequently failed to induce glomerular pathology. In addition to the *Trpc6*^*E896K*^ mutation described here, neither the *Actn4*^*K256E*^,[81, 82] nor the *Inf2*^*R218Q*^,[52] mutations induce a renal phenotype when present in heterozygous form in mice. Homozygous *Actn4*^*K255E/K255E*^ animals, though, do develop severe glomerular pathology.[81] And both heterozygous and homozygous *Inf2* mutant mice show enhanced sensitivity to injury in several injury models.[52, 56] In the case of *Trpc6*, the homozygous knock-in animals show only a transient, and slightly, higher degree of albuminuria in the albumin overload and angiotensin II infusion models. The *Trpc6*^*E896K*^ mutation, unlike the *Inf2*^*R218Q*^ mutation,[56] does not induce susceptibility to PAN. Of note, transgenic overexpression of *Trpc6* mutants, including E896K, in podocytes induces only minimal albuminuria in mice, which is not significantly different from that seen upon overexpression of wild-type *Trpc6*.[32] The reason for the relative lack of a renal phenotype in the *Trpc6*^*E896K/E896K*^ animals remains unclear. The E896K mutation is a gain-of-function mutant, as we have separately demonstrated in platelets.[44] There is a report of differences in *Trpc6* mRNA expression between mouse strains,[33] and strain-specific susceptibility to kidney, and glomerular, injury are well known.[55, 83-85] It is certainly plausible that a different genetic background, or environmental insult, is necessary to elicit the pathologic effects of *Trpc6* mutations. Alternatively, it is possible that glomerular or renal TRPC6 expression differs between humans and mice, and accounts for our findings. Review of single cell RNA expression data speaks to this possibility.[70, 86, 87] And although we did not examine TRPC6 expression in human samples, we were unable to detect the protein in murine glomeruli, or detect a GSK-inducible, Fura-2 measurable, calcium influx in murine glomerular outgrowths. Future studies comparing glomerular TRPC6 expression in different mammalian species could prove informative.

In summary, introducing a gain-of-function mutation, corresponding to a human FSGS disease mutation, into murine *Trpc6* fails to recapitulate the human glomerular disease pathology. Homozygous *Trpc6*^*E896K/E896K*^ mice do demonstrate transiently, and mildly, higher albuminuria in the albumin overload and angiotensin II infusion models, but do not display a predilection for PAN or increased interstitial fibrosis after recovery from folate-induced AKI. It remains unclear if *Trpc6* mutations require an as yet unidentified environmental or genetic hit to induce glomerular disease in mice, or if mice are intrinsically not suitable to model TRPC6-mediated human FSGS.

## Acknowledgements

We would like to thank the Beth Israel Deaconess Medical Center Histology and Transgenic Cores, and the Harvard ICCB-Longwood Screening Core, for technical assistance and expertise. We appreciated the support of the UAB O’Brien Center Core in providing serum creatinine measurements. We are indebted to Suzanne Burke for assistance with mouse surgeries, and to Dr. Martin Pollak and members of the Pollak lab for thoughtful discussions and comments.

Research reported in this publication was supported by the National Institute of Diabetes And Digestive And Kidney Diseases of the National Institutes of Health under Award Numbers R01DK115438 (to J.S.S.) and T32DK007726 (to B.R.S), and by the Klarman Scholarship Award (to J.S.S).

## References

1. D’Agati VD, Kaskel FJ, Falk RJ. Focal segmental glomerulosclerosis. The New England journal of medicine. 2011;365(25):2398–411. PubMed PMID: 22187987.

2. De Vriese AS, Wetzels JF, Glassock RJ, Sethi S, Fervenza FC. Therapeutic trials in adult FSGS: lessons learned and the road forward. Nature reviews Nephrology. 2021;17(9):619–30. doi 10.1038/s41581-021-00427-1. PubMed PMID: 34017116; PubMed Central PMCID: PMCPMC8136112.

3. Kopp JB, Anders HJ, Susztak K, Podesta MA, Remuzzi G, Hildebrandt F, et al. Podocytopathies. Nat Rev Dis Primers. 2020;6(1):68. doi: 10.1038/s41572-020-0196-7. PubMed PMID: 32792490; PubMed Central PMCID: PMCPMC8162925.

4. Sadowski CE, Lovric S, Ashraf S, Pabst WL, Gee HY, Kohl S, et al. A single-gene cause in 29.5% of cases of steroid-resistant nephrotic syndrome. J Am Soc Nephrol. 2015;26(6):1279–89. doi: 10.1681/ASN.2014050489. PubMed PMID: 25349199; PubMed Central PMCID: PMCPMC4446877.

5. Trautmann A, Bodria M, Ozaltin F, Gheisari A, Melk A, Azocar M, et al. Spectrum of steroid-resistant and congenital nephrotic syndrome in children: the PodoNet registry cohort. Clinical journal of the American Society of Nephrology : CJASN. 2015;10(4):592–600. doi: 10.2215/CJN.06260614. PubMed PMID: 25635037; PubMed Central PMCID: PMCPMC4386250.

6. Bierzynska A, McCarthy HJ, Soderquest K, Sen ES, Colby E, Ding WY, et al. Genomic and clinical profiling of a national nephrotic syndrome cohort advocates a precision medicine approach to disease management. Kidney international. 2017;91(4):937–47. doi: 10.1016/j.kint.2016.10.013. PubMed PMID: 28117080.

7. Wang M, Chun J, Genovese G, Knob AU, Benjamin A, Wilkins MS, et al. Contributions of Rare Gene Variants to Familial and Sporadic FSGS. J Am Soc Nephrol. 2019;30(9):1625–40. doi: 10.1681/ASN.2019020152. PubMed PMID: 31308072; PubMed Central PMCID: PMCPMC6727251.

8. Yao T, Udwan K, John R, Rana A, Haghighi A, Xu L, et al. Integration of Genetic Testing and Pathology for the Diagnosis of Adults with FSGS. Clinical journal of the American Society of Nephrology : CJASN. 2019;14(2):213–23. doi: 10.2215/CJN.08750718. PubMed PMID: 30647093; PubMed Central PMCID: PMCPMC6390925.

9. Groopman E, Goldstein D, Gharavi A. Diagnostic Utility of Exome Sequencing for Kidney Disease. Reply. The New England journal of medicine. 2019;380(21):2080–1. doi: 10.1056/NEJMc1903250. PubMed PMID: 31116939.

10. Li AS, Ingham JF, Lennon R. Genetic Disorders of the Glomerular Filtration Barrier. Clinical journal of the American Society of Nephrology : CJASN. 2020;15(12):1818–28. doi: 10.2215/CJN.11440919. PubMed PMID: 32205319; PubMed Central PMCID: PMCPMC7769017.

11. Hall G, Wang L, Spurney RF. TRPC Channels in Proteinuric Kidney Diseases. Cells. 2019;9(1). doi: 10.3390/cells9010044. PubMed PMID: 31877991; PubMed Central PMCID: PMCPMC7016871.

12. Reiser J, Polu KR, Moller CC, Kenlan P, Altintas MM, Wei C, et al. TRPC6 is a glomerular slit diaphragm-associated channel required for normal renal function. Nature genetics. 2005;37(7):739–44. PubMed PMID: 15924139.

13. Winn MP, Conlon PJ, Lynn KL, Farrington MK, Creazzo T, Hawkins AF, et al. A mutation in the TRPC6 cation channel causes familial focal segmental glomerulosclerosis. Science (New York, NY. 2005;308(5729):1801-4. Epub 2005/05/10. doi: 10.1126/science.1106215. PubMed PMID: 15879175.

14. Heeringa SF, Moller CC, D. J, Yue L, Hinkes B, Chernin G, et al. A novel TRPC6 mutation that causes childhood FSGS. PloS one. 2009;4(11):e7771. PubMed PMID: 19936226.

15. Mottl AK, Lu M, Fine CA, Weck KE. A novel TRPC6 mutation in a family with podocytopathy and clinical variability. BMC nephrology. 2013;14:104. Epub 2013/05/15. doi: 10.1186/1471-2369-14-104. PubMed PMID: 23663351; PubMed Central PMCID: PMC3662586.

16. Riehle M, Buscher AK, Gohlke BO, Kassmann M, Kolatsi-Joannou M, Brasen JH, et al. TRPC6 G757D Loss-of-Function Mutation Associates with FSGS. J Am Soc Nephrol. 2016. Epub 2016/02/20. doi: 10.1681/ASN.2015030318. PubMed PMID: 26892346.

17. Gigante M, Caridi G, Montemurno E, Soccio M, d’Apolito M, Cerullo G, et al. TRPC6 mutations in children with steroid-resistant nephrotic syndrome and atypical phenotype. Clinical journal of the American Society of Nephrology : CJASN. 2011;6(7):1626-34. Epub 2011/07/08. doi: 10.2215/CJN.07830910. PubMed PMID: 21734084.

18. Ramsey IS, Delling M, Clapham DE. An introduction to TRP channels. Annual review of physiology. 2006;68:619–47. PubMed PMID: 16460286.

19. Nilius B, Voets T. TRP channels: a TR(I)P through a world of multifunctional cation channels. Pflugers Arch. 2005;451(1):1-10. Epub 2005/07/14. doi: 10.1007/s00424-005-1462-y. PubMed PMID: 16012814.

20. Hofmann T, Obukhov AG, Schaefer M, Harteneck C, Gudermann T, Schultz G. Direct activation of human TRPC6 and TRPC3 channels by diacylglycerol. Nature. 1999;397(6716):259–63. PubMed PMID: 9930701.

21. Wang H, Cheng X, Tian J, Xiao Y, Tian T, Xu F, et al. TRPC channels: Structure, function, regulation and recent advances in small molecular probes. Pharmacol Ther. 2020;209:107497. doi: 10.1016/j.pharmthera.2020.107497. PubMed PMID: 32004513; PubMed Central PMCID: PMCPMC7183440.

22. Hofstra JM, Lainez S, van Kuijk WH, Schoots J, Baltissen MP, Hoefsloot LH, et al. New TRPC6 gain-of-function mutation in a non-consanguineous Dutch family with late-onset focal segmental glomerulosclerosis. Nephrology, dialysis, transplantation : official publication of the European Dialysis and Transplant Association - European Renal Association. 2013;28(7):1830–8. Epub 2013/01/08. doi: 10.1093/ndt/gfs572. PubMed PMID: 23291369.

23. Azumaya CM, Sierra-Valdez F, Cordero-Morales JF, Nakagawa T. Cryo-EM structure of the cytoplasmic domain of murine transient receptor potential cation channel subfamily C member 6 (TRPC6). The Journal of biological chemistry. 2018;293(26):10381-91. Epub 2018/05/13. doi: 10.1074/jbc.RA118.003183. PubMed PMID: 29752403; PubMed Central PMCID: PMC6028952.

24. Tang Q, Guo W, Zheng L, Wu JX, Liu M, Zhou X, et al. Structure of the receptor-activated human TRPC6 and TRPC3 ion channels. Cell research. 2018;28(7):746–55. Epub 2018/04/28. doi: 10.1038/s41422-018-0038-2. PubMed PMID: 29700422; PubMed Central PMCID: PMC6028632.

25. Bai Y, Yu X, Chen H, Horne D, White R, Wu X, et al. Structural basis for pharmacological modulation of the TRPC6 channel. eLife. 2020;9. doi: 10.7554/eLife.53311. PubMed PMID: 32149605; PubMed Central PMCID: PMCPMC7082128.

26. Guo W, Tang Q, Wei M, Kang Y, Wu JX, Chen L. Structural mechanism of human TRPC3 and TRPC6 channel regulation by their intracellular calcium-binding sites. Neuron. 2022. doi: 10.1016/j.neuron.2021.12.023. PubMed PMID: 35051376.

27. Schlondorff J, Del Camino D, Carrasquillo R, Lacey V, Pollak MR. TRPC6 mutations associated with focal segmental glomerulosclerosis cause constitutive activation of NFAT-dependent transcription. American journal of physiology Cell physiology. 2009;296(3):C558–69. PubMed PMID: 19129465.

28. Chiluiza D, Krishna S, Schumacher VA, Schlondorff J. Gain-of-function mutations in transient receptor potential C6 (TRPC6) activate extracellular signal-regulated kinases 1/2 (ERK1/2). The Journal of biological chemistry. 2013;288(25):18407–20. PubMed PMID: 23645677.

29. Yu H, Kistler A, Faridi MH, Meyer JO, Tryniszewska B, Mehta D, et al. Synaptopodin Limits TRPC6 Podocyte Surface Expression and Attenuates Proteinuria. J Am Soc Nephrol. 2016. Epub 2016/03/30. doi: 10.1681/ASN.2015080896. PubMed PMID: 27020855.

30. Talbot BE, Vandorpe DH, Stotter BR, Alper SL, Schlondorff JS. Transmembrane insertases and N-glycosylation critically determine synthesis, trafficking, and activity of the nonselective cation channel TRPC6. The Journal of biological chemistry. 2019;294(34):12655–69. doi: 10.1074/jbc.RA119.008299. PubMed PMID: 31266804; PubMed Central PMCID: PMCPMC6709635.

31. Ilatovskaya DV, Staruschenko A. TRPC6 channel as an emerging determinant of the podocyte injury susceptibility in kidney diseases. American journal of physiology. 2015;309(5):F393–7. Epub 2015/06/19. doi: 10.1152/ajprenal.00186.2015. PubMed PMID: 26084930; PubMed Central PMCID: PMC4556891.

32. Krall P, Canales CP, Kairath P, Carmona-Mora P, Molina J, Carpio JD, et al. Podocyte-specific overexpression of wild type or mutant trpc6 in mice is sufficient to cause glomerular disease. PloS one. 2010;5(9):e12859. Epub 2010/09/30. doi: 10.1371/journal.pone.0012859. PubMed PMID: 20877463; PubMed Central PMCID: PMC2942896.

33. Kunert-Keil C, Bisping F, Kruger J, Brinkmeier H. Tissue-specific expression of TRP channel genes in the mouse and its variation in three different mouse strains. BMC Genomics. 2006;7:159. PubMed PMID: 16787531.

34. Riccio A, Medhurst AD, Mattei C, Kelsell RE, Calver AR, Randall AD, et al. mRNA distribution analysis of human TRPC family in CNS and peripheral tissues. Brain research Molecular brain research. 2002;109(1-2):95–104. Epub 2003/01/18. PubMed PMID: 12531519.

35. Wu YL, Xie J, An SW, Oliver N, Barrezueta NX, Lin MH, et al. Inhibition of TRPC6 channels ameliorates renal fibrosis and contributes to renal protection by soluble klotho. Kidney international. 2017;91(4):830–41. Epub 2016/12/17. doi: 10.1016/j.kint.2016.09.039. PubMed PMID: 27979597; PubMed Central PMCID: PMC5357448.

36. Moller CC, Wei C, Altintas MM, Li J, Greka A, Ohse T, et al. Induction of TRPC6 channel in acquired forms of proteinuric kidney disease. J Am Soc Nephrol. 2007;18(1):29–36. PubMed PMID: 17167110.

37. Wang L, Jirka G, Rosenberg PB, Buckley AF, Gomez JA, Fields TA, et al. Gq signaling causes glomerular injury by activating TRPC6. The Journal of clinical investigation. 2015;125(5):1913–26. Epub 2015/04/07. doi: 10.1172/JCI76767. PubMed PMID: 25844902; PubMed Central PMCID: PMC4463190.

38. Zhang X, Song Z, Guo Y, Zhou M. The novel role of TRPC6 in vitamin D ameliorating podocyte injury in STZ-induced diabetic rats. Mol Cell Biochem. 2015;399(1-2):155-65. doi: 10.1007/s11010-014-2242-9. PubMed PMID: 25292315.

39. Eckel J, Lavin PJ, Finch EA, Mukerji N, Burch J, Gbadegesin R, et al. TRPC6 enhances angiotensin II-induced albuminuria. J Am Soc Nephrol. 2011;22(3):526–35. Epub 2011/01/25. doi: 10.1681/ASN.2010050522. PubMed PMID: 21258036; PubMed Central PMCID: PMC3060446.

40. Spires D, Ilatovskaya DV, Levchenko V, North PE, Geurts AM, Palygin O, et al. Protective role of Trpc6 knockout in the progression of diabetic kidney disease. American journal of physiology. 2018;315(4):F1091–F7. Epub 2018/06/21. doi: 10.1152/ajprenal.00155.2018. PubMed PMID: 29923767.

41. Kim EY, Yazdizadeh Shotorbani P, Dryer SE. Trpc6 inactivation confers protection in a model of severe nephrosis in rats. J Mol Med (Berl). 2018;96(7):631–44. doi: 10.1007/s00109-018-1648-3. PubMed PMID: 29785489; PubMed Central PMCID: PMCPMC6015123.

42. Kistler AD, Singh G, Altintas MM, Yu H, Fernandez IC, Gu C, et al. Transient receptor potential channel 6 (TRPC6) protects podocytes during complement-mediated glomerular disease. The Journal of biological chemistry. 2013;288(51):36598–609. Epub 2013/11/07. doi: 10.1074/jbc.M113.488122. PubMed PMID: 24194522; PubMed Central PMCID: PMC3868772.

43. Wang L, Chang JH, Buckley AF, Spurney RF. Knockout of TRPC6 promotes insulin resistance and exacerbates glomerular injury in Akita mice. Kidney international. 2019;95(2):321–32. Epub 2019/01/23. doi: 10.1016/j.kint.2018.09.026. PubMed PMID: 30665571.

44. Boekell KL, Brown BJ, Talbot BE, Schlondorff JS. Trpc6 gain-of-function disease mutation enhances phosphatidylserine exposure in murine platelets. bioRxiv. 2022:2022.02.01.478727. doi: 10.1101/2022.02.01.478727.

45. Dietrich A, Mederos YSM, Gollasch M, Gross V, Storch U, Dubrovska G, et al. Increased vascular smooth muscle contractility in TRPC6-/-mice. Molecular and cellular biology. 2005;25(16):6980–9. PubMed PMID: 16055711.

46. Hartleben B, Godel M, Meyer-Schwesinger C, Liu S, Ulrich T, Kobler S, et al. Autophagy influences glomerular disease susceptibility and maintains podocyte homeostasis in aging mice. The Journal of clinical investigation. 2010;120(4):1084–96. Epub 2010/03/05. doi: 10.1172/JCI39492. PubMed PMID: 20200449; PubMed Central PMCID: PMC2846040.

47. Refaeli I, Hughes MR, Wong AK, Bissonnette MLZ, Roskelley CD, Wayne Vogl A, et al. Distinct Functional Requirements for Podocalyxin in Immature and Mature Podocytes Reveal Mechanisms of Human Kidney Disease. Sci Rep. 2020;10(1):9419. doi: 10.1038/s41598-020-64907-3. PubMed PMID: 32523052; PubMed Central PMCID: PMCPMC7286918.

48. Stotter BR, Talbot BE, Capen DE, Artelt N, Zeng J, Matsumoto Y, et al. Cosmc-dependent mucin-type O-linked glycosylation is essential for podocyte function. American journal of physiology. 2020;318(2):F518–F30. doi: 10.1152/ajprenal.00399.2019. PubMed PMID: 31904283; PubMed Central PMCID: PMCPMC7052656.

49. Street JM, Souza AC, Alvarez-Prats A, Horino T, Hu X, Yuen PS, et al. Automated quantification of renal fibrosis with Sirius Red and polarization contrast microscopy. Physiol Rep. 2014;2(7). doi: 10.14814/phy2.12088. PubMed PMID: 25052492; PubMed Central PMCID: PMCPMC4187565.

50. Im JH, Muschel JJ. Protocol for Murine/Mouse Platelets Isolation and Their Reintroduction in vivo. Bio-protocol. 2017;7(4):e2132. doi: 10.21769/BioProtoc.2132.

51. Takemoto M, Asker N, Gerhardt H, Lundkvist A, Johansson BR, Saito Y, et al. A new method for large scale isolation of kidney glomeruli from mice. The American journal of pathology. 2002;161(3):799–805. doi: 10.1016/S0002-9440(10)64239-3. PubMed PMID: 12213707; PubMed Central PMCID: PMCPMC1867262.

52. Subramanian B, Sun H, Yan P, Charoonratana VT, Higgs HN, Wang F, et al. Mice with mutant Inf2 show impaired podocyte and slit diaphragm integrity in response to protamine-induced kidney injury. Kidney international. 2016;90(2):363–72. Epub 2016/06/29. doi: 10.1016/j.kint.2016.04.020. PubMed PMID: 27350175.

53. Yumura W, Sugino N, Nagasawa R, Kubo S, Hirokawa K, Maruyama N. Age-associated changes in renal glomeruli of mice. Exp Gerontol. 1989;24(3):237–49. doi: 10.1016/0531-5565(89)90015-6. PubMed PMID: 2731581.

54. Zheng F, Plati AR, Potier M, Schulman Y, Berho M, Banerjee A, et al. Resistance to glomerulosclerosis in B6 mice disappears after menopause. The American journal of pathology. 2003;162(4):1339–48. doi: 10.1016/S0002-9440(10)63929-6. PubMed PMID: 12651625; PubMed Central PMCID: PMCPMC1851217.

55. Ishola DA, Jr., van der Giezen DM, Hahnel B, Goldschmeding R, Kriz W, Koomans HA, et al. In mice, proteinuria and renal inflammatory responses to albumin overload are strain-dependent. Nephrology, dialysis, transplantation : official publication of the European Dialysis and Transplant Association -European Renal Association. 2006;21(3):591–7. Epub 2005/12/06. doi: 10.1093/ndt/gfi303. PubMed PMID: 16326737.

56. Sun H, Perez-Gill C, Schlondorff JS, Subramanian B, Pollak MR. Dysregulated Dynein-Mediated Trafficking of Nephrin Causes INF2-related Podocytopathy. J Am Soc Nephrol. 2021;32(2):307–22. doi: 10.1681/ASN.2020081109. PubMed PMID: 33443052; PubMed Central PMCID: PMCPMC8054882.

57. Onohara N, Nishida M, Inoue R, Kobayashi H, Sumimoto H, Sato Y, et al. TRPC3 and TRPC6 are essential for angiotensin II-induced cardiac hypertrophy. The EMBO journal. 2006;25(22):5305–16. PubMed PMID: 17082763.

58. Shi J, Ju M, Saleh SN, Albert AP, Large WA. TRPC6 channels stimulated by angiotensin II are inhibited by TRPC1/C5 channel activity through a Ca2+- and PKC-dependent mechanism in native vascular myocytes. The Journal of physiology. 2010;588(Pt 19):3671-82. doi: 10.1113/jphysiol.2010.194621. PubMed PMID: 20660561; PubMed Central PMCID: PMCPMC2998219.

59. Tian D, Jacobo SM, Billing D, Rozkalne A, Gage SD, Anagnostou T, et al. Antagonistic regulation of actin dynamics and cell motility by TRPC5 and TRPC6 channels. Science signaling. 2010;3(145):ra77. PubMed PMID: 20978238.

60. Ilatovskaya DV, Palygin O, Chubinskiy-Nadezhdin V, Negulyaev YA, Ma R, Birnbaumer L, et al. Angiotensin II has acute effects on TRPC6 channels in podocytes of freshly isolated glomeruli. Kidney international. 2014;86(3):506–14. Epub 2014/03/22. doi: 10.1038/ki.2014.71. PubMed PMID: 24646854; PubMed Central PMCID: PMC4149864.

61. Anderson M, Roshanravan H, Khine J, Dryer SE. Angiotensin II activation of TRPC6 channels in rat podocytes requires generation of reactive oxygen species. Journal of cellular physiology. 2014;229(4):434–42. Epub 2013/09/17. doi: 10.1002/jcp.24461. PubMed PMID: 24037962.

62. Imgrund M, Grone E, Grone HJ, Kretzler M, Holzman L, Schlondorff D, et al. Re-expression of the developmental gene Pax-2 during experimental acute tubular necrosis in mice 1. Kidney international. 1999;56(4):1423–31. doi: 10.1046/j.1523-1755.1999.00663.x. PubMed PMID: 10504494.

63. Long DA, Woolf AS, Suda T, Yuan HT. Increased renal angiopoietin-1 expression in folic acid-induced nephrotoxicity in mice. J Am Soc Nephrol. 2001;12(12):2721–31. doi: 10.1681/ASN.V12122721. PubMed PMID: 11729241.

64. Yang HC, Zuo Y, Fogo AB. Models of chronic kidney disease. Drug Discov Today Dis Models. 2010;7(1-2):13-9. doi: 10.1016/j.ddmod.2010.08.002. PubMed PMID: 21286234; PubMed Central PMCID: PMCPMC3030258.

65. Zhang Y, Yin N, Sun A, Wu Q, Hu W, Hou X, et al. Transient Receptor Potential Channel 6 Knockout Ameliorates Kidney Fibrosis by Inhibition of Epithelial-Mesenchymal Transition. Front Cell Dev Biol. 2020;8:602703. doi: 10.3389/fcell.2020.602703. PubMed PMID: 33520986; PubMed Central PMCID: PMCPMC7843578.

66. Lin BL, Matera D, Doerner JF, Zheng N, Del Camino D, Mishra S, et al. In vivo selective inhibition of TRPC6 by antagonist BI 749327 ameliorates fibrosis and dysfunction in cardiac and renal disease. Proceedings of the National Academy of Sciences of the United States of America. 2019;116(20):10156–61. doi: 10.1073/pnas.1815354116. PubMed PMID: 31028142; PubMed Central PMCID: PMCPMC6525474.

67. Kong W, Haschler TN, Nurnberg B, Kramer S, Gollasch M, Marko L. Renal Fibrosis, Immune Cell Infiltration and Changes of TRPC Channel Expression after Unilateral Ureteral Obstruction in Trpc6-/-Mice. Cell Physiol Biochem. 2019;52(6):1484–502. doi: 10.33594/000000103. PubMed PMID: 31099508.

68. Chung HS, Kim GE, Holewinski RJ, Venkatraman V, Zhu G, Bedja D, et al. Transient receptor potential channel 6 regulates abnormal cardiac S-nitrosylation in Duchenne muscular dystrophy. Proceedings of the National Academy of Sciences of the United States of America. 2017;114(50):E10763–E71. doi: 10.1073/pnas.1712623114. PubMed PMID: 29187535; PubMed Central PMCID: PMCPMC5740634.

69. Hofmann K, Fiedler S, Vierkotten S, Weber J, Klee S, Jia J, et al. Classical transient receptor potential 6 (TRPC6) channels support myofibroblast differentiation and development of experimental pulmonary fibrosis. Biochimica et biophysica acta Molecular basis of disease. 2017;1863(2):560–8. Epub 2016/12/10. doi: 10.1016/j.bbadis.2016.12.002. PubMed PMID: 27932059.

70. Chung JJ, Goldstein L, Chen YJ, Lee J, Webster JD, Roose-Girma M, et al. Single-Cell Transcriptome Profiling of the Kidney Glomerulus Identifies Key Cell Types and Reactions to Injury. J Am Soc Nephrol. 2020;31(10):2341–54. doi: 10.1681/ASN.2020020220. PubMed PMID: 32651223; PubMed Central PMCID: PMCPMC7609001.

71. Doleschal B, Primessnig U, Wolkart G, Wolf S, Schernthaner M, Lichtenegger M, et al. TRPC3 contributes to regulation of cardiac contractility and arrhythmogenesis by dynamic interaction with NCX1. Cardiovascular research. 2015;106(1):163–73. Epub 2015/01/30. doi: 10.1093/cvr/cvv022. PubMed PMID: 25631581; PubMed Central PMCID: PMC4362401.

72. Xu X, Lozinskaya I, Costell M, Lin Z, Ball JA, Bernard R, et al. Characterization of Small Molecule TRPC3 and TRPC6 agonist and antagonists. Biophysical journal. 2013;104(2):454a.

73. Fischer KG, Saueressig U, Jacobshagen C, Wichelmann A, Pavenstadt H. Extracellular nucleotides regulate cellular functions of podocytes in culture. American journal of physiology. 2001;281(6):F1075–81. doi: 10.1152/ajprenal.2001.281.6.F1075. PubMed PMID: 11704558.

74. Forst AL, Olteanu VS, Mollet G, Wlodkowski T, Schaefer F, Dietrich A, et al. Podocyte Purinergic P2X4 Channels Are Mechanotransducers That Mediate Cytoskeletal Disorganization. J Am Soc Nephrol. 2016;27(3):848–62. doi: 10.1681/ASN.2014111144. PubMed PMID: 26160898; PubMed Central PMCID: PMCPMC4769195.

75. Roshanravan H, Dryer SE. ATP acting through P2Y receptors causes activation of podocyte TRPC6 channels: role of podocin and reactive oxygen species. American journal of physiology. 2014;306(9):F1088–97. Epub 2014/02/21. doi: 10.1152/ajprenal.00661.2013. PubMed PMID: 24553432.

76. Palygin O, Klemens CA, Isaeva E, Levchenko V, Spires DR, Dissanayake LV, et al. Characterization of purinergic receptor 2 signaling in podocytes from diabetic kidneys. iScience. 2021;24(6):102528. doi: 10.1016/j.isci.2021.102528. PubMed PMID: 34142040; PubMed Central PMCID: PMCPMC8188476.

77. Guan Y, Nakano D, Zhang Y, Li L, Liu W, Nishida M, et al. A protease-activated receptor-1 antagonist protects against podocyte injury in a mouse model of nephropathy. J Pharmacol Sci. 2017. doi: 10.1016/j.jphs.2017.09.002. PubMed PMID: 29110957.

78. Taal MW, Brenner BM. Renoprotective benefits of RAS inhibition: from ACEI to angiotensin II antagonists. Kidney international. 2000;57(5):1803–17. doi: 10.1046/j.1523-1755.2000.00031.x. PubMed PMID: 10792600.

79. Kurahara LH, Sumiyoshi M, Aoyagi K, Hiraishi K, Nakajima K, Nakagawa M, et al. Intestinal myofibroblast TRPC6 channel may contribute to stenotic fibrosis in Crohn’s disease. Inflamm Bowel Dis. 2015;21(3):496–506. doi: 10.1097/MIB.0000000000000295. PubMed PMID: 25647156.

80. Kim EY, Dryer SE. Effects of TRPC6 Inactivation on Glomerulosclerosis and Renal Fibrosis in Aging Rats. Cells. 2021;10(4). doi: 10.3390/cells10040856. PubMed PMID: 33918778; PubMed Central PMCID: PMCPMC8070418.

81. Yao J, Le TC, Kos CH, Henderson JM, Allen PG, Denker BM, et al. Alpha-actinin-4-mediated FSGS: an inherited kidney disease caused by an aggregated and rapidly degraded cytoskeletal protein. PLoS biology. 2004;2(6):e167. Epub 2004/06/23. doi: 10.1371/journal.pbio.0020167. PubMed PMID: 15208719; PubMed Central PMCID: PMC423141.

82. Henderson JM, Al-Waheeb S, Weins A, Dandapani SV, Pollak MR. Mice with altered alpha-actinin-4 expression have distinct morphologic patterns of glomerular disease. Kidney international. 2008;73(6):741–50. Epub 2008/01/11. doi: 10.1038/sj.ki.5002751. PubMed PMID: 18185509; PubMed Central PMCID: PMC2980842.

83. Tsaih SW, Pezzolesi MG, Yuan R, Warram JH, Krolewski AS, Korstanje R. Genetic analysis of albuminuria in aging mice and concordance with loci for human diabetic nephropathy found in a genome-wide association scan. Kidney international. 2010;77(3):201–10. doi: 10.1038/ki.2009.434. PubMed PMID: 19924099; PubMed Central PMCID: PMCPMC2807478.

84. Pippin JW, Brinkkoetter PT, Cormack-Aboud FC, Durvasula RV, Hauser PV, Kowalewska J, et al. Inducible rodent models of acquired podocyte diseases. American journal of physiology. 2009;296(2):F213–29. doi: 10.1152/ajprenal.90421.2008. PubMed PMID: 18784259.

85. de Mik SM, Hoogduijn MJ, de Bruin RW, Dor FJ. Pathophysiology and treatment of focal segmental glomerulosclerosis: the role of animal models. BMC nephrology. 2013;14:74. doi: 10.1186/1471-2369-14-74. PubMed PMID: 23547922; PubMed Central PMCID: PMCPMC3637050.

86. Muto Y, Wilson PC, Ledru N, Wu H, Dimke H, Waikar SS, et al. Single cell transcriptional and chromatin accessibility profiling redefine cellular heterogeneity in the adult human kidney. Nature communications. 2021;12(1):2190. doi: 10.1038/s41467-021-22368-w. PubMed PMID: 33850129; PubMed Central PMCID: PMCPMC8044133.

87. Kirita Y, Wu H, Uchimura K, Wilson PC, Humphreys BD. Cell profiling of mouse acute kidney injury reveals conserved cellular responses to injury. Proceedings of the National Academy of Sciences of the United States of America. 2020;117(27):15874–83. doi: 10.1073/pnas.2005477117. PubMed PMID: 32571916; PubMed Central PMCID: PMCPMC7355049.

